# Inter-brain functional connectivity: Are we measuring the right thing?

**DOI:** 10.1101/2025.07.02.662718

**Authors:** Juan Camilo Avendano-Diaz, Patrick Sothmann, Riitta Hari, Lauri Parkkonen

**Affiliations:** Department of Neuroscience and Biomedical Engineering, Aalto University, Espoo, Finland; Department of Art and Media, Aalto University, Espoo, Finland

**Keywords:** Hyperscanning, Inter-brain functional connectivity, Brain oscillations, Signal-to-noise ratio (SNR), Social interaction

## Abstract

Hyperscanning—the simultaneous recording of brain activity from multiple individuals— and the study of inter-brain synchronization is gaining popularity in social neuroscience. MEG/EEG hyperscanning studies often estimate inter-brain functional connectivity using phase-based metrics applied to oscillatory brain signals, assuming matching peak frequencies between the individuals studied. However, in reality, peak frequencies typically differ between subjects and between brain regions. Using simulated MEG/EEG signals, we systematically assessed how inter-individual frequency differences affect commonly used connectivity measures. Phase-based metrics were highly sensitive to frequency differences across individuals, whereas amplitude envelope correlation remained robust, offering more reliable connectivity estimates. Our results underscore the need for connectivity metrics specifically tailored to inter-brain analyses. These findings are relevant to a range of disciplines that are increasingly integrating hyperscanning into their methodological toolkits.

## Introduction

Research on the brain basis of social interaction is complementing the traditional brain imaging studies with more naturalistic and interactive approaches. This methodological shift has been driven by calls to study two or more interacting individuals simultaneously, enabling insights that cannot be obtained through single-person paradigms (1–3). As a result, hyperscanning, the simultaneous measurement of brain activity of two or more individuals (4), has been increasingly adopted across various noninvasive neuroimaging modalities, as reflected in the number of published hyperscanning studies during the last decade (5) (see Fig.1A).

**Fig. 1.**
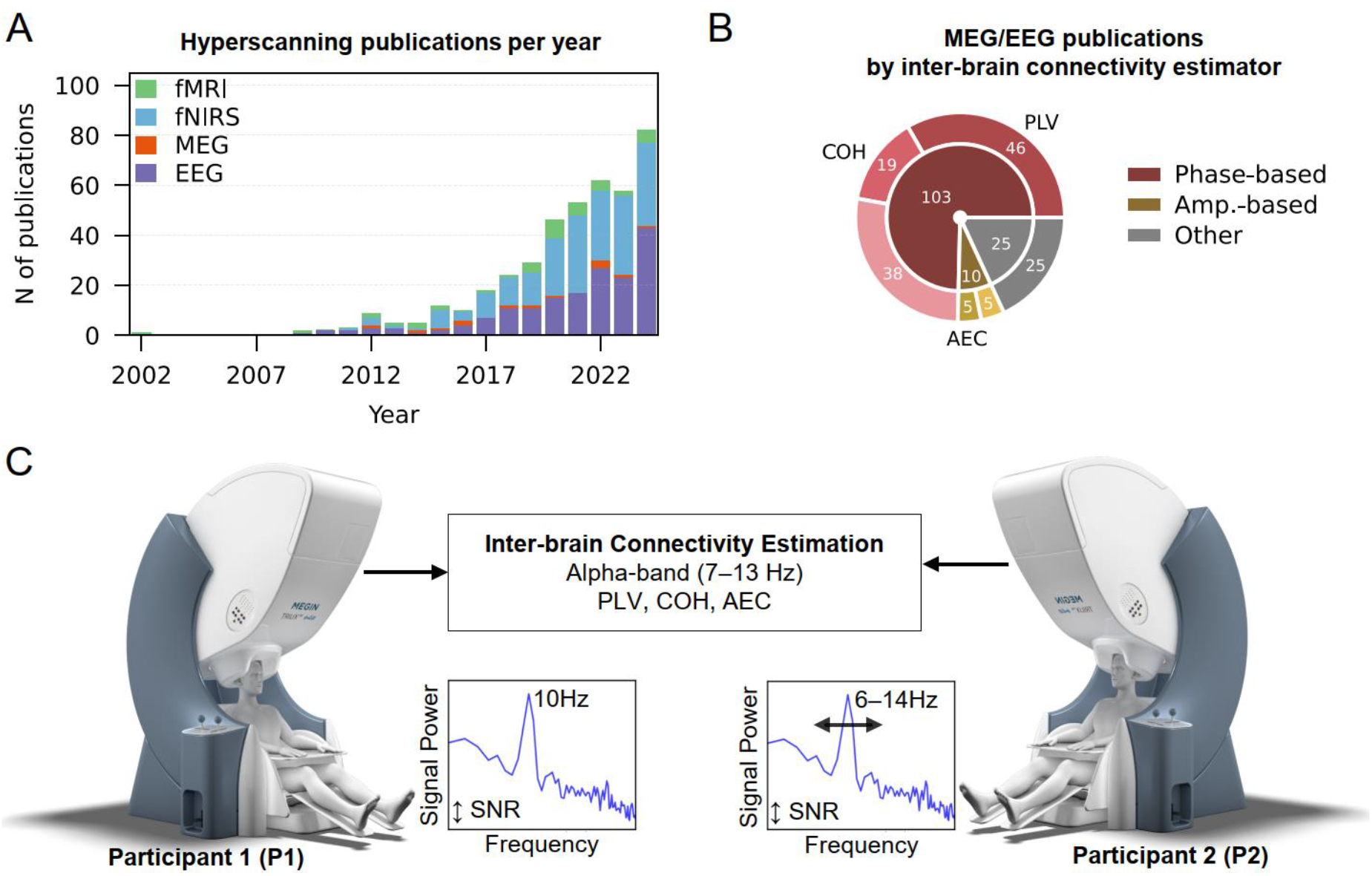
Hyperscanning landscape and simulation setup for evaluating inter-brain functional connectivity. (A) The number of hyperscanning publications per year, grouped by neuroimaging modality. (B) MEG/EEG hyperscanning publications estimating inter-brain coupling, grouped by the type of connectivity metric employed (Phase-based, Amplitude-based, and Other approaches). The number of publications that use phase-locking value (PLV), coherence (COH), and amplitude-envelope correlation (AEC) is highlighted. The data were retrieved from Scopus using relevant search terms. (C) Schematic of our simulation framework. We generated MEG/EEG signals for two participants P1 and P2. The oscillatory signals P1 had a fixed frequency of 10 Hz. For P2, a set of oscillatory signals with varying frequency (6–14 Hz, 0.1-Hz steps) was created to simulate inter-individual variability in oscillatory peak frequencies. We estimated inter-brain functional connectivity in the alpha band (7–13 Hz), using PLV, COH, and AEC. Connectivity estimations were repeated while varying the signal-to-noise ratio (SNR) levels. MEG systems image source: Megin Oy, 2024, https://megin.com/what-is-meg/

Hyperscanning studies typically estimate interdependencies between the brain signals of interacting individuals using functional connectivity metrics (5). The core assumption is that synchronization of brain activity between individuals, also referred to as inter-brain coupling, reflects aspects of social interaction (6). However, most electroencephalography (EEG) and magnetoencephalography (MEG) hyperscanning studies rely on phase-based connectivity metrics originally developed to quantify frequency-specific synchronization between signals within a single brain (7) (see Fig.1B). These metrics assess dependencies between oscillations at equal frequencies. Yet, spontaneous neural oscillations vary in their peak frequencies between individuals, and within individuals across brain areas and tasks (8, 9). Even within certain brain regions, the sources of the predominant oscillations can change dynamically. For instance, the sources of a spindle of the posterior alpha rhythm typically remain stable for less than a second (10). Moreover, signal mixing across brain regions, an issue that affects more EEG than MEG data, can also contribute to inter-individual frequency differences. As a result, due to unaccounted inter-individual frequency differences, many MEG/EEG hyperscanning studies may have misestimated the functional interdependencies across interacting brains and their potential relation to social interaction.

We simulated oscillatory MEG/EEG signals for two participants (P1 and P2; see Fig.1C) to systematically investigate how inter-individual differences in oscillatory peak frequency affect estimates of inter-brain functional connectivity. We computed connectivity in the 7–13 Hz “alpha band” using commonly employed phase-based metrics (see Fig.1B)—phase-locking value (PLV) and coherence (COH)—and evaluated amplitude envelope correlation (AEC) as an alternative. We also examined how signal-to-noise ratio (SNR) affects these estimates.

## Results

Fig. 2 summarizes the results. Estimates are referenced to the situation when P1 and P2 share a peak oscillatory frequency of 10 Hz.

**Fig. 2.**
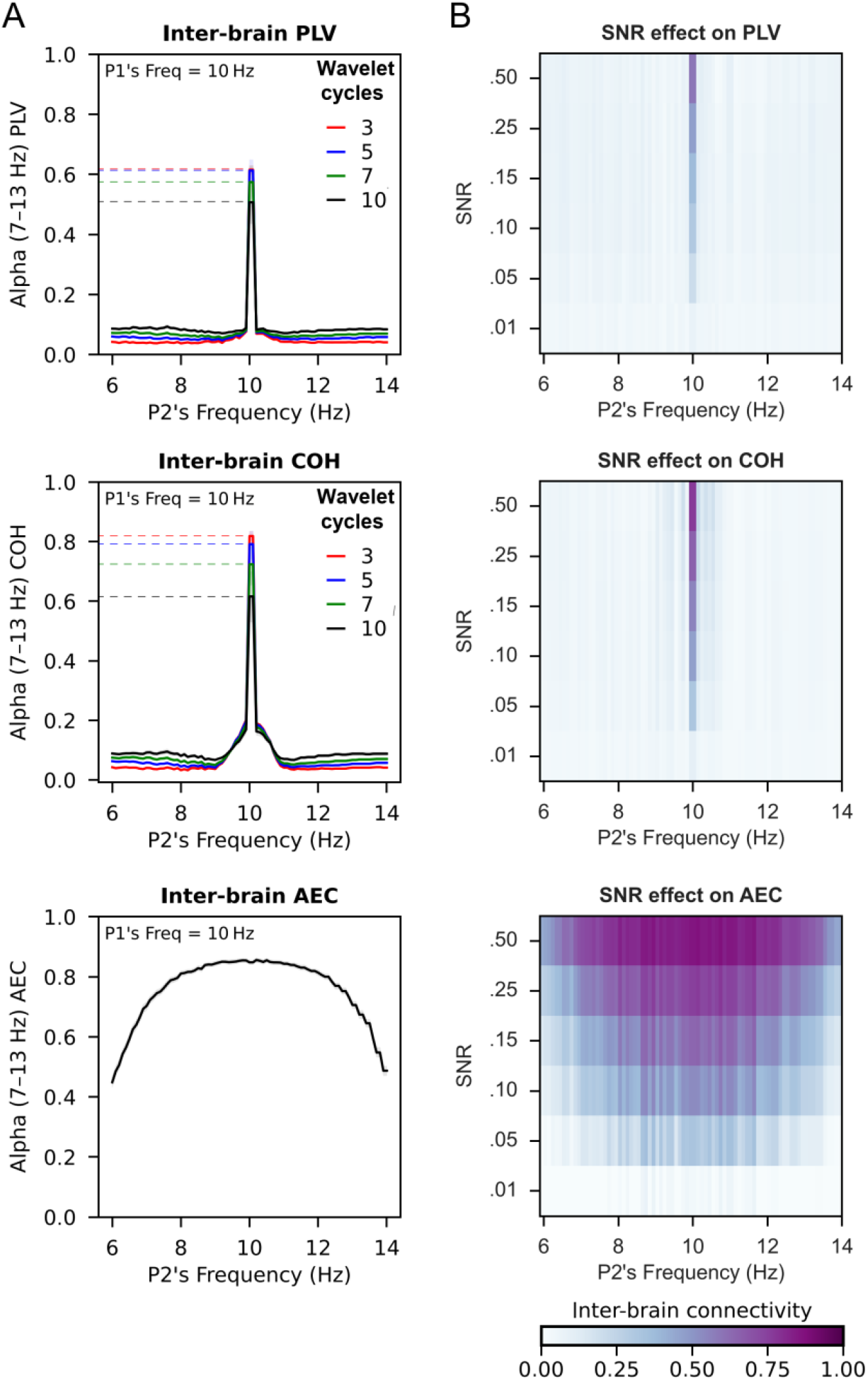
Effect of oscillatory peak frequency differences and signal-to-noise ratio on estimates of inter-brain functional connectivity. (A) Inter-brain connectivity between Participant 1 (P1; fixed frequency of 10 Hz) and Participant 2 (P2; 6–14 Hz, 0.1Hz steps), estimated using PLV (top), COH (middle), and AEC (bottom) in the alpha band (7–13 Hz). The horizontal axis shows the peak frequencies of P2’s signals. Shaded areas represent 95% confidence intervals. (B) Inter-brain connectivity estimates across SNR levels. For PLV and COH, results are shown for 5 wavelet cycles; similar patterns were observed for 3, 7, and 10 cycles. Overall, phase-based connectivity estimates, unlike AEC, declined rapidly as P2’s frequency diverged from P1’s (10 Hz), regardless of the number of wavelet cycles employed.

### Phase-based inter-brain connectivity

PLV and COH were highly sensitive to inter-individual peak frequency differences (see Fig. 2A) so that the accuracy of the estimates of inter-brain connectivity declined rapidly as the peak frequencies of P1 and P2 diverged. This effect was consistently observed across all applied numbers of wavelet cycles (3, 5, 7, and 10).

### Amplitude-envelope-based inter-brain connectivity

In contrast to phase-based metrics, the AEC results were more robust to differences between the peak frequencies of P1 and P2 (see Fig. 2A, bottom panel). The inter-brain connectivity estimates remained sensitive and stable across the frequency range of interest.

### SNR-dependence of inter-brain connectivity

Even when the peak oscillatory frequencies of P1 and P2 were equal (10 Hz), SNR reduction decreased connectivity estimates for all metrics (see Fig. 2B). Notably, when the peak frequencies of P1 and P2 differed, the frequency mismatch dominated the results, overriding the influence of SNR.

## Discussion

Our findings highlight a critical methodological limitation in the MEG/EEG hyperscanning literature: the phase-based connectivity metrics commonly used to examine inter-brain synchrony, here represented by PLV and COH, are highly sensitive to inter-individual differences in the peak frequency of spontaneous brain oscillations. Our simulations show that even small differences between the peak frequencies of participants in a hyperscanning study can lead to substantial misestimations of inter-brain connectivity based on this oscillatory activity, which could compromise subsequent interpretations and potential links to social interaction.

Problems of using phase-based connectivity metrics in hyperscanning have been discussed earlier, demonstrating for example that PLV and COH are prone to detecting spurious couplings even in the absence of true inter-brain synchronization (11), and that common arbitrary methodological decisions can bias inter-brain coupling estimates (12). Furthermore, in studies of infant–adult interactions, where large frequency mismatches are unavoidable, it has been acknowledged that employing conventional phase-based metrics may result in misleading interpretations (13). Alternative metrics that do not require strict frequency alignment, including cross-frequency coupling and symbolic mutual information, might help address these limitations (13). However, the validity and interpretability of these approaches in hyperscanning still warrant systematic evaluation.

In contrast to phase-based metrics, our findings showed that AEC remained robust to peak frequency mismatch. This (expected) robustness makes AEC potentially more suitable for hyperscanning analyses. It is important to note that we simulated MEG/EEG signals from both individuals as continuous sinusoidal oscillations amplitude-modulated by an identical aperiodic signal, which is an idealized scenario for AEC that is unlikely to occur in real life. Moreover, AEC has its own limitations. For instance, it can be influenced by non-neural signal correlations, such as shared noise or stimulus-locked co-modulation, and typically requires longer data segments to yield reliable estimates (5, 14). As with any connectivity measure, its appropriateness depends on the specific research question, and the experimental context (14).

Our simulations further showed that the accuracy of inter-brain connectivity estimates was modulated by the SNR. These findings add to previous calls to carefully consider SNR differences between experimental conditions when interpreting results of intra- and inter-brain connectivity (9, 12, 14). Our findings also suggest that the effect of inter-individual peak frequency differences has a dominant influence on the accuracy of inter-brain connectivity estimates compared to that of the SNR.

In sum, while hyperscanning offers a promising window into the brain basis of social interaction, it also presents unique methodological challenges (3). Careful consideration of the limitations and validity of current connectivity metrics in inter-brain research is essential for obtaining reliable and interpretable insights into how the brain activity of interacting individuals may support smooth social interaction. More broadly, our findings highlight the need to develop or adopt inter-brain connectivity metrics specifically tuned for hyperscanning datasets. Or, alternatively, to temper the enthusiasm for summarizing the richness and complexity of two-brain data into simplified indices of inter-brain synchronization.

Moving forward, we encourage the hyperscanning field to expand its methodological repertoire by developing and systematically comparing connectivity metrics that take human physiology into account (e.g., metrics that are robust to inter-individual variability in oscillatory properties). To establish the validity and relevance of these metrics, it is crucial to relate inter-brain coupling measures to behavioral, affective, psychological, and contextual factors of social interaction. We also advocate for greater replication efforts and the joint analysis of intra- and inter-brain connectivity, which remain rare but could illuminate the unique contributions of shared and individual brain processes. Such steps are essential to ensure that hyperscanning continues to evolve into a rigorous approach for investigating the brain basis of real-world social interactions.

## Materials and Methods

We simulated MEG/EEG signals from two individuals (P1 and P2; see Fig.1C), systematically varying P2’s peak frequency (6–14 Hz, 0.1-Hz steps), while keeping P1’s frequency fixed at 10 Hz. We simulated these signals as the sum of three components: (1) an oscillation at the desired frequency, amplitude-modulated by an aperiodic 1/f^χ^ signal (0.005–0.5 Hz, χ = 0.5) in line with human physiology (15), (2) an aperiodic component with a pink noise exponent (χ = 1), and (3) white noise. For each scenario, we created with NeuroDSP (16) 50 trials of 30 s each, at a sampling rate of 1000 Hz. We estimated inter-brain functional connectivity between signals representing P1 and P2, in the alpha frequency band (7–13 Hz), using PLV, COH, and AEC. For PLV and COH, spectral decomposition was performed using complex Morlet wavelets with 3, 5, 7, and 10 cycles. AEC was obtained by applying the Hilbert transform to the bandpass-filtered signals to extract amplitude envelopes. Connectivity estimates were computed over time for each trial and then averaged across the 50 simulations. To manipulate SNR, we adjusted the amplitude of both P1 and P2’s signals while keeping the noise level constant, following the procedure used in Ref. (9). We then assessed the effect of SNR on inter-brain connectivity using the connectivity estimation framework described above. We used MNE-Connectivity (17) for connectivity estimation.

## Acknowledgments

We acknowledge the computational resources provided by the Aalto Science-IT project. J.C.A-D. and L.P. were supported by Business Finland (grant “DIGIMIND” 7981/31/2022). J.C.A-D., R.H., and L.P. were supported by the Norman Loveless Memorial Fund.

## Notes

### Competing Interest Statement

The authors declare the following competing interests: L.P. has a part-time employment with the MEG device vendor Megin Oy. The other authors declare no competing interests.

